# A machine learning strategy that leverages large datasets to boost statistical power in small-scale experiments

**DOI:** 10.1101/849331

**Authors:** William E. Fondrie, William S. Noble

## Abstract

Machine learning methods have proven invaluable for increasing the sensitivity of peptide detection in proteomics experiments. Most modern tools, such as Percolator and PeptideProphet, use semi-supervised algorithms to learn models directly from the datasets that they analyze. Although these methods are effective for many proteomics experiments, we suspected that they may be suboptimal for experiments of smaller scale. In this work, we found that the power and consistency of Percolator results was reduced as the size of the experiment was decreased. As an alternative, we propose a different operating mode for Percolator: learn a model with Percolator from a large dataset and use the learned model to evaluate the small-scale experiment. We call this a “static modeling” approach, in contrast to Percolator’s usual “dynamic model” that is trained anew for each dataset. We applied this static modeling approach to two settings: small, gel-based experiments and single-cell proteomics. In both cases, static models increased the yield of detected peptides and eliminated the model-induced variability of the standard dynamic approach. These results suggest that static models are a powerful tool for bringing the full benefits of Percolator and other semi-supervised algorithms to small-scale experiments.

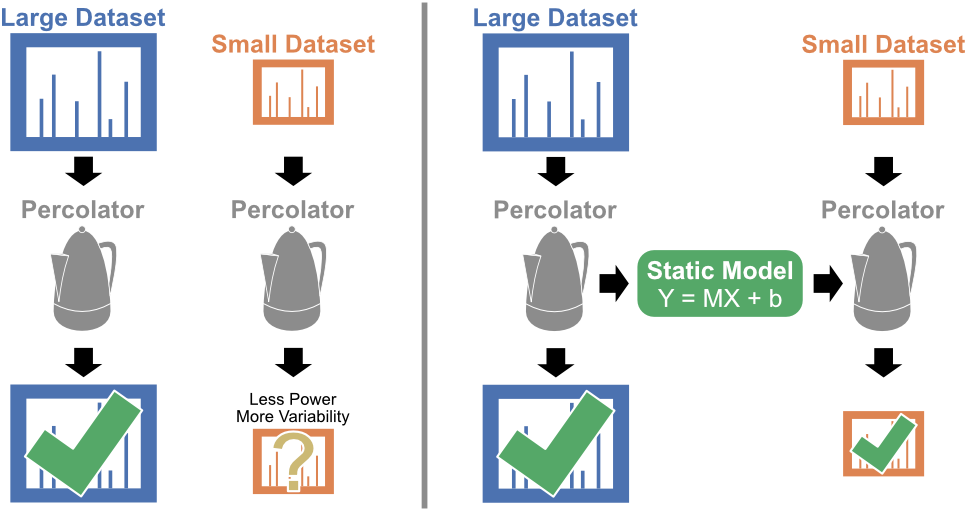

## 1 Introduction

The assignment of peptide sequences to tandem mass spectra is a fundamental task in any proteomics experiment [1]. This task is most often performed by a database search algorithm, which was first introduced with the SEQUEST search engine in 1994 [2]. Database search algorithms score the theoretical mass spectra of peptides from a selected sequence database against the acquired mass spectra from the experiment, yielding a set of peptide-spectrum matches (PSMs). Although the score functions of individual search engines may differ greatly, the scores reported by each are intended to reflect the quality of PSMs.

Machine learning strategies to re-score PSMs were first introduced due to their ability to integrate multiple, orthogonal scores and features from database search engines, thereby improving the sensitivity of peptide detection. Two such methods were proposed simultaneously: one based on linear discriminant analysis [3] and another that used support vector machine (SVM) models to a similar end [4]. Critically, both methods were examples of supervised learning: they relied on fully labeled datasets—where the correct and incorrect PSMs could be determined *a priori*—to train their respective models. These trained models were then used to predict a new score for the PSMs of a new dataset. We refer to the models used by these methods as static, meaning their parameters do not change when the dataset is changed. Critically, the success of these static models depended on the quality of the labeled dataset that was used for training and how well it reflected the datasets that were subsequently analyzed. The original version of PeptideProphet [3] attempted to mitigate the latter weakness by using an expectation-maximization algorithm to learn a mapping function between the discriminant score and the estimated probabilities in an unsupervised fashion from the new dataset.

The advent of target-decoy competition [5] as a method to estimate the error rates of mass spectrum identification also brought forth the opportunity for improvements in machine learning strategies. The Percolator algorithm is a semi-supervised learning method to re-score the PSMs of a proteomics experiment without relying on a separate, fully-labeled dataset for training [6]. Percolator attempts to discriminate correct PSMs from incorrect PSMs by defining positive PSMs as those originating from target peptide sequences below a specified false-discovery rate (FDR) threshold and negative PSMs as those arising from decoy peptide sequences. Starting with positive PSMs accepted using the best starting feature or an initial set of weights, Percolator iteratively accumulates positive PSMs by fitting SVM models to the set of positive and negative (decoy) PSMs. The set of positive PSMs is then updated using the model predictions from each iteration. Percolator performs this model training in a three-fold cross-validation scheme, such that the new score for each PSM is produced by a model that did not use that PSM for training; this strategy avoids Percolator overfitting to the PSMs that it is attempting to re-score [7]. Percolator’s dynamic model training allows the algorithm to adapt to the unique qualities of each dataset. Since its introduction, Percolator has become a popular option to post-process database search results from a variety of search engines, including SEQUEST [2], Comet [8], Tide [9], Mascot [10], MS-GF+ [11], and X! Tandem [12]. In all these cases, Percolator demonstrably improves the power to detect peptides from tandem mass spectra and provides a statistical framework to interpret the results [13]. A recent, independent comparison of post-processing methods across three search engines and eight datasets found that “combinations involving Percolator achieved markedly more peptide and protein identifications at the same FDR level than the other 12 combinations for all data sets.” [14]

Percolator was originally designed for the typical scale of proteomics experiments, where thousands of confident PSMs are expected to be found. However, some experiments naturally result in data of smaller scale, such as applications that require low abundance samples as input. We hypothesize that analyzing small-scale experiments with Percolator leads to decreased sensitivity and increased variability in the resulting PSMs and peptides when compared to Percolator’s performance on experiments of larger scale. We verified this hypothesis empirically by systematically evaluating the performance of Percolator when analyzing experiments of decreasing scale. Consequently, we propose the use of static Percolator models—models learned by Percolator on an external dataset—to analyze small-scale experiments, and we demonstrate the potential of the method on two applications: the analysis of small gel-based experiments and for single-cell proteomics. Support for static models is now available in Percolator, both in the stand-alone version (http://percolator.ms) and as part of the Crux toolkit (http://crux.ms) [15].

## 2 Methods

### 2.1 Datasets

We analyzed three previously described proteomics datasets, collected using data-dependent acquisition methods. Each of the datasets was searched using the Tide search engine [9] implemented in the Crux mass spectrometry analysis toolkit [15]. In all cases, we employed the combined p-value score function [16]. We selected the precursor *m*/*z* window using Param-Medic [17] with a fallback value of 50 ppm, and we set the fragment ion tolerance to 0.02 Da. The protein databases (described below) were processed using Tide to generate a shuffled decoy peptide sequence for each target peptide sequence in the database, while preserving both terminal amino acids. The unique aspects of each dataset, including a brief description, the protein database, and the modifications we used, are described below. We chose these parameters to best reflect those from the original analysis of each dataset.

#### 2.1.1 A draft human proteome

The Kim *et al.* draft map of the human proteome dataset [18] consists of approximately 25 million mass spectra from more than 2,000 mass spectrometry acquisitions using LTQ-Orbitrap Elite and Velos mass spectrometers. The raw data files were downloaded from PRIDE Archive [19] (ID: PXD000561) and converted to ms2 format using msconvert [20] with peak-picking and deisotoping filters (“peakPicking vendor msLevel=2” and “MS2Deisotope Poisson”).

We searched the ms2 files against the canonical UniProt [21] human proteome (71,785 protein sequences, downloaded February 17, 2016), allowing for two missed cleavages and the following variable modifications: protein N-terminal acetylation and oxidation of methionine. Carbamidomethylation of cysteine was specified as a static modification, and the peptides were limited to two variable modifications. These parameters resulted in a database of 8,684,473 peptides.

#### 2.1.2 Histone gel bands

The Basilicata *et al.* histone dataset [22] consists of 94 mass spectrometry acquisitions, each analyzing a histone-enriched SDS-PAGE gel band from various patient derived fibroblasts using a Q-Exactive mass spectrometer. The raw data files were downloaded from PRIDE Archive (ID: PXD009317) and converted to mzML format using ThermoRawFileParser [23], with vendor peak-picking enabled.

We searched the mzML files against the canonical UniProt/Swiss-Prot human proteome (20,416 protein sequences, downloaded September 6, 2019), allowing for two missed cleavages. In addition to the modifications allowed for the Kim *et al.* dataset, we allowed the following variable modifications with a maximum of two variable modifications per peptide, as used in the original publication: deamidation of asparagine and glutamine, methylation of lysine and arginine, and lysine acetylation, trimethylation, propionylation, and propionyl-methylation. These parameters resulted in a large database of 218,942,133 peptides.

#### 2.1.3 Single-cell proteomics by mass spectrometry (SCoPE-MS) experiments

The Specht *et al.* single-cell proteomics dataset [24] consists of two distinct sets of experiments: a quality control dataset consisting of 76 mass spectrometry acquisitions and a macrophage differentiation dataset consisting of 69 mass spectrometry acquisitions, each acquired using a Q-Exactive mass spectrometer. In both cases, each acquisition corresponds to a single experiment analyzing multiple single cells using the tandem mass tag (TMT) 10-plex reagents as described in the original paper [24]. The raw data files were downloaded from MassIVE [25] (ID: MSV000083945) and converted to mzML format using ThermoRawFileParser, with vendor peak-picking enabled.

We searched the mzML files against the canonical UniProt/Swiss-Prot human proteome (20,416 protein sequences, downloaded September 6, 2019), allowing for two missed cleavages. We included the TMT 10-plex modification of lysine and the peptide N-terminus as a static modification, but carbamidomethylation of cysteine was not included. Additionally, we considered the oxidation of methionine, protein N-terminal acetylation, and deamidation of asparagine as variable modifications. These parameters resulted in a database of 21,160,904 peptides. Four of the macrophage differentiation experiments resulted in no PSMs at a 1% FDR threshold and were excluded from further analysis.

### 2.2 Downsampling to simulate small-scale experiments

We sampled sets of PSMs from the human proteome dataset to simulate experiments of decreasing size analyzed with Percolator. We first uniformly sampled 100,000 PSMs from the 23,330,311 total PSMs to be held out as a common test set for evaluation. From the remaining PSMs in the dataset, we uniformly sampled 1,000,000 PSMs to serve as the base training set.

We postulate that small datasets can be defined in two ways: either the dataset has few PSMs in total, or the dataset has few confident PSMs. In order to simulate both of these conditions, we downsampled the base training set to smaller training sets by two methods. To simulate the condition of few total PSMs, we created incrementally smaller datasets by uniformly sampling PSMs from the base training set, independently. For the condition where few confident PSMs are present, we defined confident PSMs as PSMs accepted at 1% FDR after the base training set was analyzed with Percolator. We uniformly sampled confident PSMs from the base training set and supplemented these PSMs with an additional sample of “unconfident” PSMs, such that the total number of PSMs was constant at 100,000.

Our goal was to investigate changes to the sensitivity and variability of the results obtained from Percolator on small datasets; hence, we analyzed each training set with Percolator using five unique random seeds. We then used the learned weights from each training set to re-score the PSMs in the test set. This allowed us to directly compare the performance of the Percolator models across the various training set sizes.

### 2.3 Comparison of static and dynamic Percolator models

We analyzed individual experiments from the histone gel band and SCoPE-MS datasets using both Percolator’s standard dynamic model training and a static model to evaluate whether static models improve the performance of Percolator on small-scale experiments. For the histone gel band dataset, we used the aggregate of 10 experiments (80,031 top-ranked PSMs, 797,291 total PSMs)—each defined by a single mass spectrometry acquisition—as a training set to generate a static Percolator model. The remaining experiments were individually analyzed with Percolator in duplicate, once using a dynamic model and once using the static model.

We performed similar analyses on the SCoPE-MS experiments. In this case, we used the 76 quality control experiments as a training set (558,279 top-ranked PSMs, 5,576,478 total PSMs) for a static Percolator model. The 64 macrophage SCoPE-MS experiments were then analyzed individually with Percolator, again using both dynamic and static models.

### 2.4 Data and code availability

Percolator is an open-source project and is publicly available on GitHub (https://github.com/percolator/percolator). All of the datasets used in this manuscript are already available through their respective ProteomeXchange [26] partner repositories. All code used for these analyses and to generate the figures of this manuscript is available on BitBucket (https://www.bitbucket.org/noblelab/static-percolator).

## 3 Results

### 3.1 Percolator loses power and exhibits increased variability with small datasets

Our first goal was to systematically test if Percolator’s performance declines when the analyzed experiments are small. Here, we use “small” to describe two possible conditions: an experiment that yielded few total PSMs or one that yielded few confident PSMs. Either condition can occur due to a wide variety of causes. The former condition may be due to a limited number of tandem mass spectra acquired—because of a short chromatographic gradient or low signal during acquisition—or when only a subset of PSMs are of interest. Likewise, the condition of few confident PSMs can occur when the acquired tandem mass spectra are of poor quality, such as when limited by the analyte abundance in the sample. In either case, we suspected that these limitations would hinder the performance of the Percolator algorithm.

To generate progressively smaller experiments to analyze with Percolator, we chose to down-sample multiple sets of PSMs from the Kim *et al.* draft human proteome dataset. We performed this downsampling in two ways: 1) by sampling decreasing numbers of total PSMs and 2) by sampling decreasing numbers of confident PSMs, while keeping the total number of PSMs constant. We analyzed each of these subsets with Percolator using five random seeds to assess the variability of the algorithm; the random seeds only serve to randomly assign PSMs to the cross-validation splits, since optimization of the SVM models is deterministic. We then assessed the models learned by Percolator by applying them to a held-out test set consisting of the 100,000 PSMs. Since this test set was constant for all of the downsampled experiments, it allowed us to compare the learned models to one another. We evaluated these experiments using the q-values estimated by Percolator for the test set PSMs, where a q-value is the minimal FDR threshold at which a given PSM is accepted.

We began our analysis by investigating pairwise comparisons of the q-values estimated for the test set PSMs using the models learned from the downsampled experiments. We first compared each of the total PSM downsampling experiments to those obtained by an experiment with 100,000 total PSMs (Figure 1 A–D). As the total number of PSMs analyzed with Percolator decreased, the resulting q-values became increasingly divergent. Furthermore, many test set PSMs would be considered confident only when the largest experiment was analyzed. We found a similar trend when we performed the analogous analysis on the confident PSM downsampling experiments (Figure 1 E–H). Again, the resulting q-values became increasingly divergent as Percolator was provided fewer confident PSMs. In addition to the pairwise comparisons between the downsampling experiments, we also investigated the reproducibility of the test set q-values when the same experiment was analyzed with Percolator using different random seeds (Figure S1). In both the cases of downsampling total or confident PSMs, the concordance between Percolator analyses with two random seeds decreases with the scale of the experiment.

**Figure 1:**
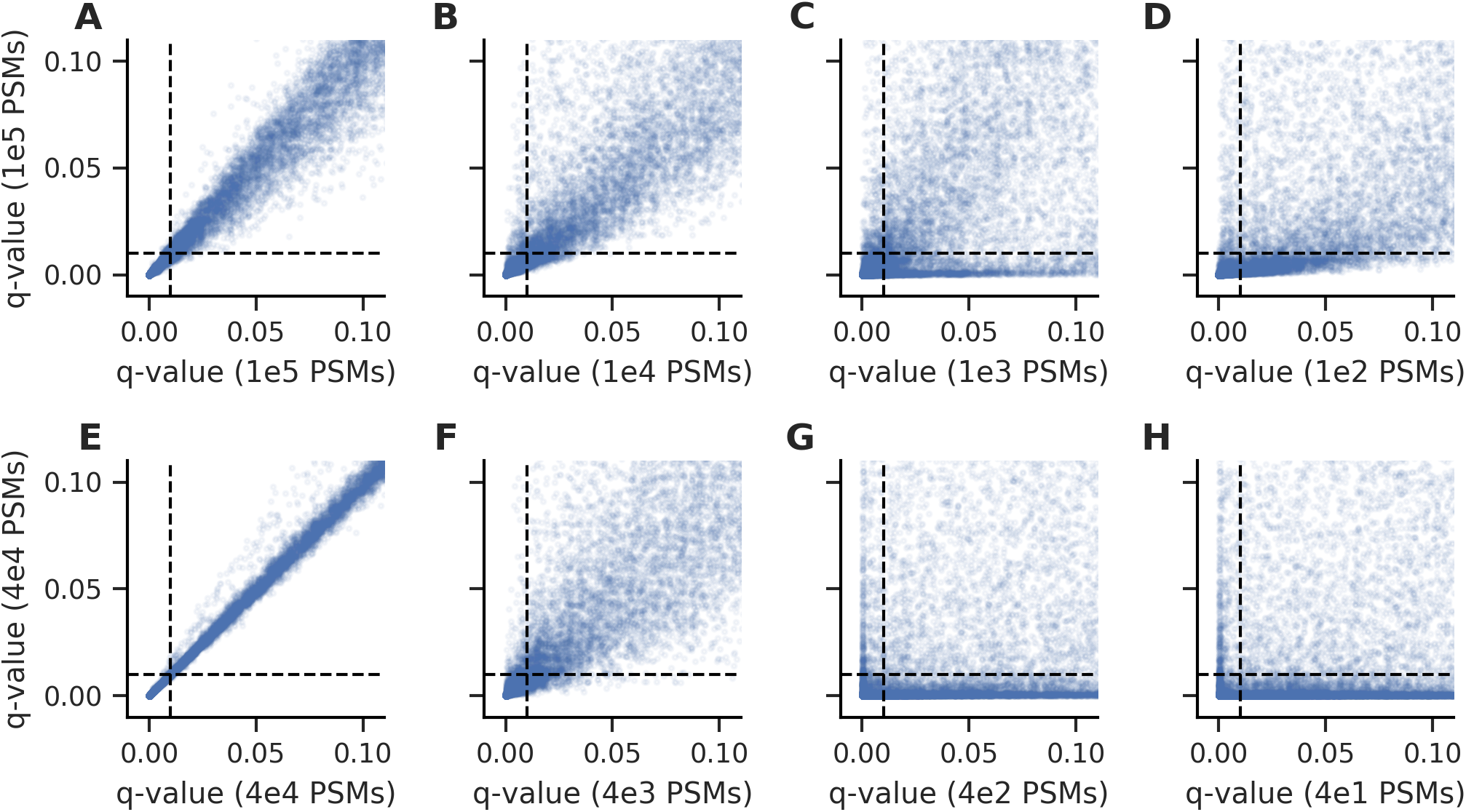
The q-values resulting from Percolator became increasingly discordant as the experiment size is decreased. The test set PSMs accepted at low q-values diverged as the total number of PSMs in the training experiments were decreased from (A) 100,000 PSMs to (B) 10,000, (C) 1,000, and (D) 100 PSMs. Likewise, the test set PSMs diverged as the number of confident PSMs in the training experiments were decreased from (E) 40,000 PSMs to (F) 4,000, (G) 400, and (H) 40 PSMs. Each point indicates a test set PSM and the dashed lines indicate a 1% FDR threshold.

In an effort to understand the overall effects of the downsampling experiments on our Percolator analyses, we looked at the number of test set PSMs that were accepted at 1% FDR by each model. As expected, a consistent decrease in the number of accepted test set PSMs was observed as the total number of PSMs analyzed with Percolator was decreased (Figure 2 A). Additionally, we saw more dramatic decreases in the number of accepted test set PSMs after downsampling confident PSMs (Figure 2 B). In both cases, the variability of the accepted test set PSMs increased when analyzing smaller experiments with Percolator, which is consistent with the results from our pairwise comparisons. Together, these experiments led us to conclude that Percolator suffers a loss of power and increased variability in its confidence estimates when used to analyze experiments of sufficiently small scale. The SVM models that Percolator trains rely on positive and negative examples to find the best separating hyperplane for a set of PSMs; hence, it is unsurprising that the resulting models may be suboptimal when there are few examples to learn from.

**Figure 2:**
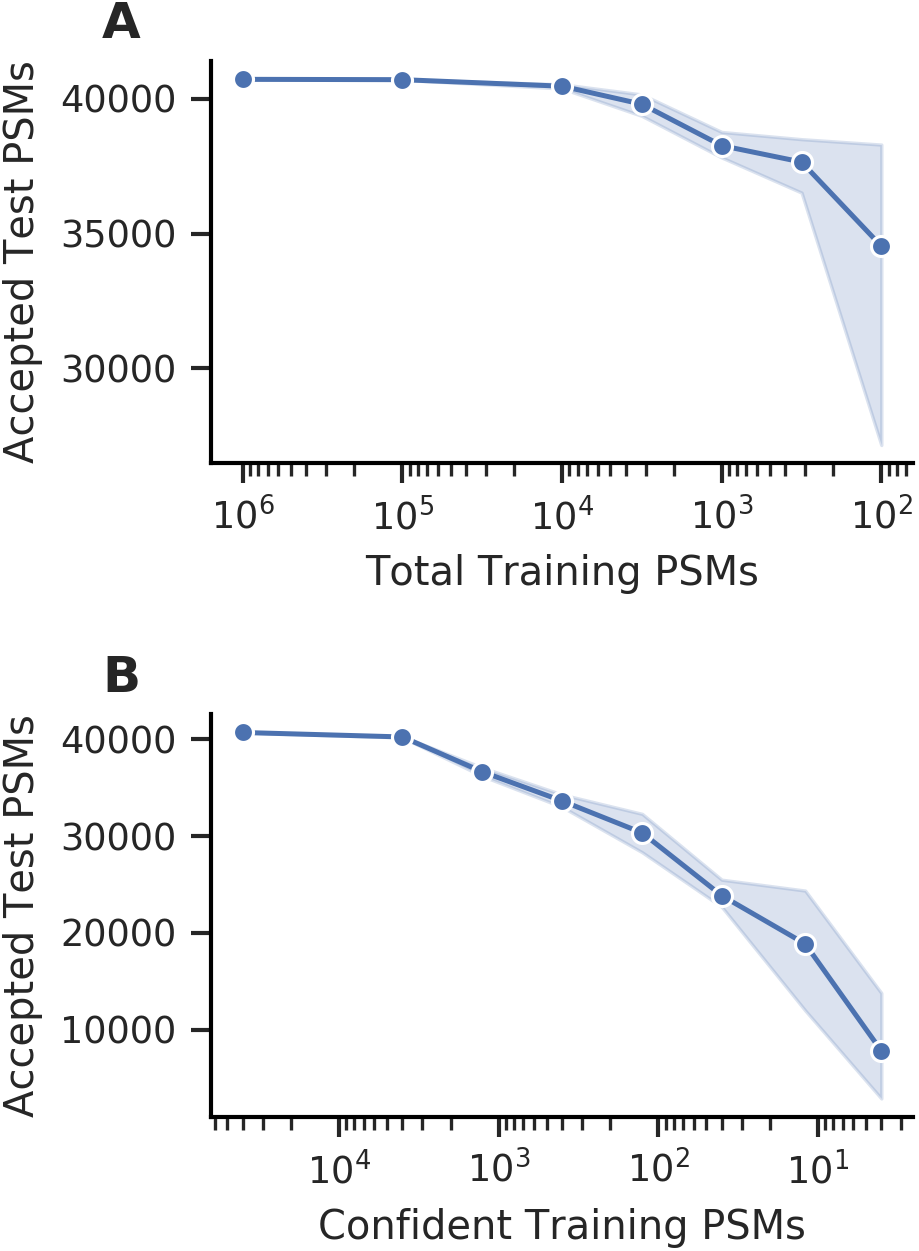
The number of accepted test set PSMs at 1% FDR decreases with smaller training experiments. (A) Decreasing the total number of training PSMs resulted in a gradual decrease in performance and increased variability. (B) Decreasing the number of confident training PSMs resulted in a rapid decline in performance. The points indicate the mean of five random seeds and the shaded region indicates the 95% confidence interval.

While we posit that most proteomics experiments are of sufficient size to use Percolator reliably, there are experiments that do not fully realize the benefits of Percolator due to their small size. In these cases, we propose the use of static Percolator models—models that were learned by Percolator from one or a collection of external experiments—to bring the benefits of Percolator to small-scale experiments. To demonstrate the effectiveness of this approach, we have applied it to two settings in the following sections: the analysis of individual SDS-PAGE gel bands and the analysis of a single-cell proteomics dataset.

### 3.2 Static models increase the peptides detected from SDS-PAGE gel bands

A routine task for proteomics laboratories and core facilities has long been the identification and characterization of the proteins from specific regions of a gel-based separation [27]. Despite the age of these gel-based methods, these types of analyses are still valuable for answering specific biological questions [28–31]. In these settings, a common goal is to detect the proteins—and potentially the post-translational modifications—that are present within an excised band from a 1-dimensional SDS-PAGE gel. We suspect that the scale of these experiments is often small, such that they would hinder the expected performance of Percolator, and we propose that the use of static Percolator models could improve the sensitivity of peptide detection in these experiments.

We sought to test these hypotheses using the collection of experiments performed by Basilicata *et al.* [22]. This dataset consists of 94 experiments each analyzing an excised SDS-PAGE gel band that is enriched for histones from a variety of patient-derived fibroblasts. We held out 10 experiments as a training set for a static Percolator model, then analyzed the remaining 84 experiments with Percolator either using its standard, dynamically trained models or the static model learned from the training set. The use of a static model increased the number of peptides detected at 1% FDR for 52 of the 72 experiments (Figure 3 A). Using the number of PSMs accepted at 1% FDR before Percolator as a proxy for the number of confident PSMs in each experiment, we found a trend that follows what was observed in the downsampling experiments—the experiments with fewer confident PSMs benefited the most from using a static model. In some cases the number of detected peptides increased by as much as 22% when using the static model (Figure 3 B). In contrast, other experiments detected fewer peptides when a static model was used; however, the largest loss in peptides detected for any experiment was only 4.5% (Figure 3 C). Furthermore, inspection of the peptide yield across low FDR thresholds for this experiment does not indicate a systematic loss of peptides. Rather, this apparent loss is well within the expected variance inherent in decoy-based FDR estimates [32, 33]. Overall these results suggest that it is useful to consider a static model for the analysis of excised gel bands, particularly when ample data is available to train a static Percolator model.

**Figure 3:**
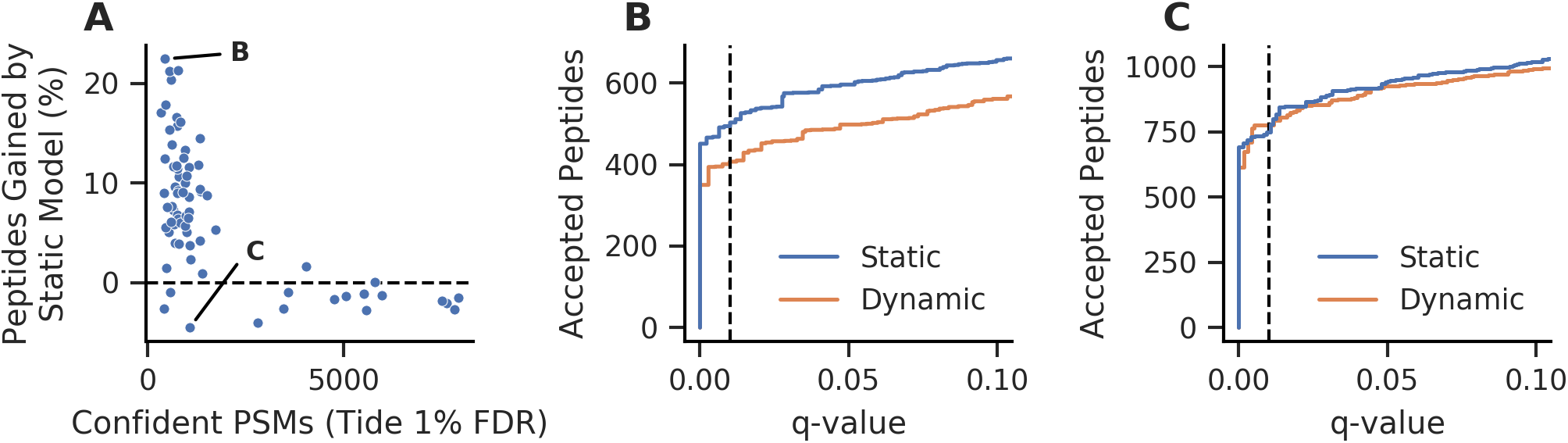
Static Percolator models improve the detection of peptides from individual histone SDS-PAGE gel bands. (A) The static Percolator model improved the sensitivity of peptide detection at 1% FDR for most experiments when compared to the Percolator’s typical dynamic model training, particularly when few confident PSMs were found. Confident PSMs were defined as PSMs accepted at 1% FDR using the Tide combined p-value before Percolator analysis. The labeled points are further investigated in panels B and C. (B) The experiment for which we observed the largest peptide gain at 1% FDR also had consistent gains across the low FDR range. (C) The experiment that resulted in the largest peptide loss at 1% FDR is inconsistent over the low FDR range. The dashed lines indicate 1% FDR.

### 3.3 Static models reduce missing peptides across SCoPE-MS experiments

While the analysis of small, gel-based experiments represents a long-standing, routine task for proteomics laboratories, the analysis of single cells is an emerging field and an exciting future for proteomics technologies [34]. Currently, several methods are making progress toward performing proteomics experiments at single-cell resolution [24, 35–37]. Critically, all of these methods inherntly rely on acquiring tandem mass spectra from samples with low analyte abundance, which in turn results in fewer acquired mass spectra and many mass spectra of insufficient quality for identification; hence, we sought to determine if static Percolator models could improve peptide detection rates in single-cell proteomics experiments.

We decided to analyze the single-cell proteomics datasets from the recent update to the SCoPE-MS method [24]. The SCoPE-MS method uses TMT to multiplex single cells for analysis, while reserving one or more TMT channels for a “carrier sample,” i.e. a sample where on the order of 100 cells are analyzed to boost the detectable signal in the mass spectra. As an isobaric labeling experiment, the relative quantitation for peptides from single cells is performed by extracting the reporter ion signals from tandem mass spectra that are confidently identified with peptides. Thus, to compare a peptide across multiple experiments the peptide must be confidently detected in each. Specht *et al.* presented two distinct sets of SCoPE-MS experiments that we have used in our analyses: a quality control dataset and a macrophage differentiation dataset, each consisting of a large collection of experiments.

For our analyses, we used the quality control SCoPE-MS dataset to train a static Percolator model. We then individually analyzed 65 experiments from the macrophage differentiation SCoPE-MS dataset with Percolator, using either the standard, dynamically trained models or the static model learned from the training set. It is worth noting that these experiments could have been analyzed jointly with Percolator, but FDR estimation would then need to be performed outside of the Percolator framework; this is because we were interested in the peptides detected in each run, rather than in aggregate. In our analyses, we found that the static model consistently yielded increased numbers of PSMs and peptides (Figure 4 A). At the PSM level, 57 of the 65 SCoPE-MS experiments (88%) had increased yield at 1% FDR, with the largest loss of only 3.2%. Similarly, 58 of 65 SCoPE-MS experiments (89%) had an increased peptide yield at 1% FDR, indicating that the static model improved the sensitivity of peptide detection in these SCoPE-MS experiments.

**Figure 4:**
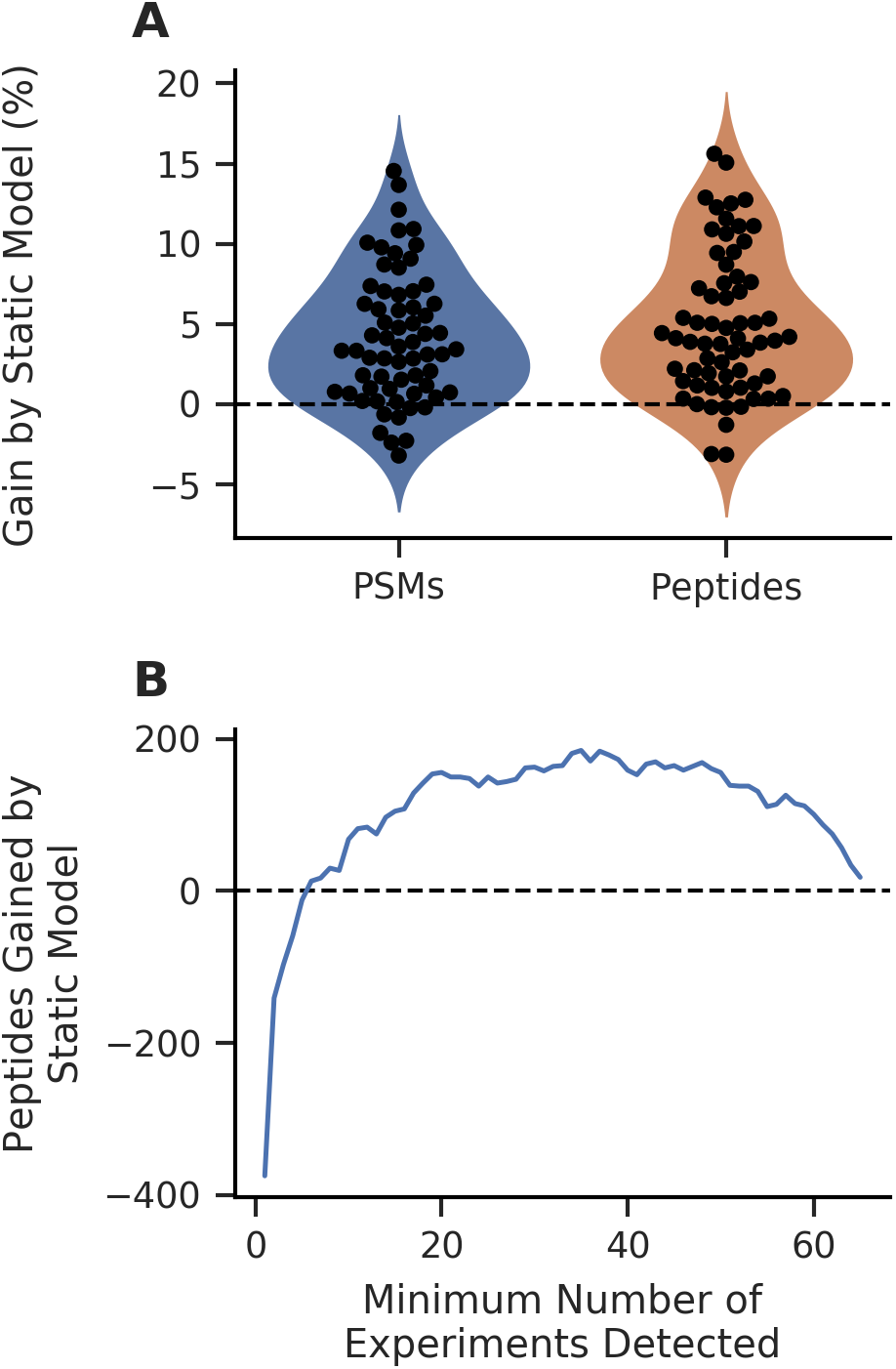
The static Percolator model improves the number of PSMs and peptides detected in SCoPE-MS experiments. (A) The static model resulted in improved sensitivity at both the PSM and peptide levels for most experiments when compared to Percolator’s dynamic model training. (B) More peptides are consistently detected across more experiments at 1% FDR when using the static model, resulting in a less sparse peptide-experiment matrix.

We argue that the number of peptides consistently detected across experiments is of greater concern than the total number of peptides detected in the individual experiments for a SCoPE-MS dataset. To investigate how peptide detection consistency is affected by the Percolator analysis with a static model, we counted the number of peptides detected at 1% FDR that were shared between at least a specified number of experiments (Figure S2 A). We observed an appreciable increase in the number of peptides that were found in large numbers of experiments using the static model, despite detecting fewer overall unique peptides (Figure 4 B); this trend started at peptides detected in at least six or more experiments. These results indicate that peptides are detected with improved consistency across experiments when the Percolator analysis is performed with a static model.

Finally, we suspected that the improved consistency of peptide detection could be only partially attributed to the increased sensitivity provided by the static Percolator model. Specifically, we postulated that the static model also resulted in a more consistent ranking of peptides across experiments. We tested this hypothesis by allowing the peptides obtained from the dynamic Percolator models to yield the same number of peptides as obtained with the static Percolator model at 1% FDR, effectively nullifying the sensitivity advantage of the static model. Comparison of the peptides detected with the static model to these expanded results from the dynamic models revealed that the static models still increase the number of peptides that are detected across many experiments (Figure S2 B). In light of these findings, we concluded that Percolator analysis with static models increased the consistency of peptide detection across SCoPE-MS experiments due to both increased sensitivity and decreased variability in peptide rankings.

## 4 Discussion

In this work, we investigated the robustness of the Percolator algorithm when it is used to analyze small-scale experiments. This investigation revealed that the results obtained from Percolator lose power and increase in variability as experiments decrease in scale. Unfortunately, the question of how small an experiment can be before the algorithm suffers cannot be explicitly answered for all cases. Rather, this will depend on a variety of experimental parameters including the search engine and features used, the diversity of the peptides analyzed, and the quality of the mass spectra that were collected. However, we would generally suggest that a minimum of 5,000 total PSMs— assuming that approximately one third of the PSMs can be assigned confidently—be used as a minimum experiment size when using the Tide search engine, based on the presented results.

As an alternative to the dynamic model training that is normally performed by Percolator, we demonstrated that the use of static models improved the consistency and statistical power to detect peptides in small-scale experiments. Although we explored only two experimental designs, we expect that there are many types of experiments that may benefit from static models due to the small scale of the results they yield. For example, studies of ancient proteins must often be performed on limited, degraded material, such as in the recent analysis of ancient enamel proteins from Early Pleistocene specimens [38]. More commonly, the scale of an experiment can be reduced by limiting Percolator analysis to a subset of the total PSMs, such as when only PSMs containing a specific modification or arising from particular proteins are of interest. In all these cases, larger datasets may be leveraged to increase the yield and consistency of peptides from Percolator using a static model.

Furthermore, one benefit we did not explore in this work was the increase in speed gained from using a static model. The static model approach requires training a model only once, prior to the analysis of the experiment of interest; hence, a static model does not need the iterative training involved the dynamic modeling approach. The increased speed that the static model approach provides will allow for models to be used in time-sensitive applications. One example of an application that could benefit from this static model approach would be real-time database searching [39]. The Orbiter method uses the Comet search engine to score PSMs in real-time, attempting to maximize the peptides detected and quantified in a TMT experiment. A static model learned by Percolator could potentially be used to increase the sensitivity of the Orbiter method by providing a more powerful score for each PSM as the spectrum is collected.

Support for static models is available as of Percolator version 3.04 (http://percolator.ms) and is also available within the Crux toolkit (http://crux.ms), both of which are open source projects.

## 5 Acknowledgments

We are grateful to Lukas Käll and Matthew The for helpful discussions on the implementation of static models in Percolator and reviewing the code for this feature. The research reported in this publication was supported by the National Institutes of Health awards T32 HG000035 and P41 GM103533.

