## Supplemental Figures for "A machine learning strategy that leverages large datasets to boost statistical power in small-scale experiments"

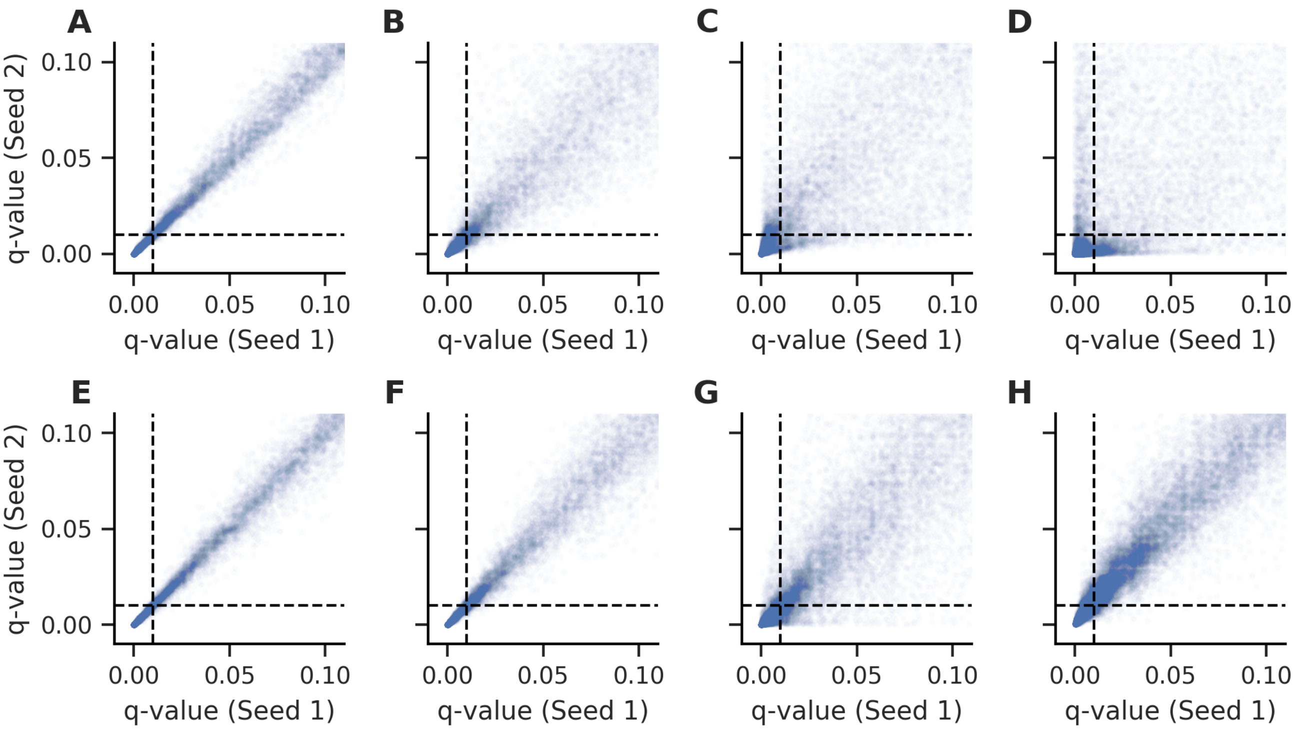


Supplementary Figure 1: The q-values resulting from different Percolator seeds become increasingly discordant as the experiment size is decreased. The test set PSMs accepted when Percolator is run with two different random seeds diverge as the total number of PSMs in the training experiment are decreased from (A) 100,000 PSMs to (B) 10,000, (C) 1,000, and (D) 100 PSMs. Similarly, the test set PSMs accepted between random seeds diverge as the number of confident PSMs in the training experiment are decreased from (E) 40,000 PSMs to (F) 4,000, (G) 400, and (H) 40 PSMs. The dashed lines indicate a q-value of 0.01.

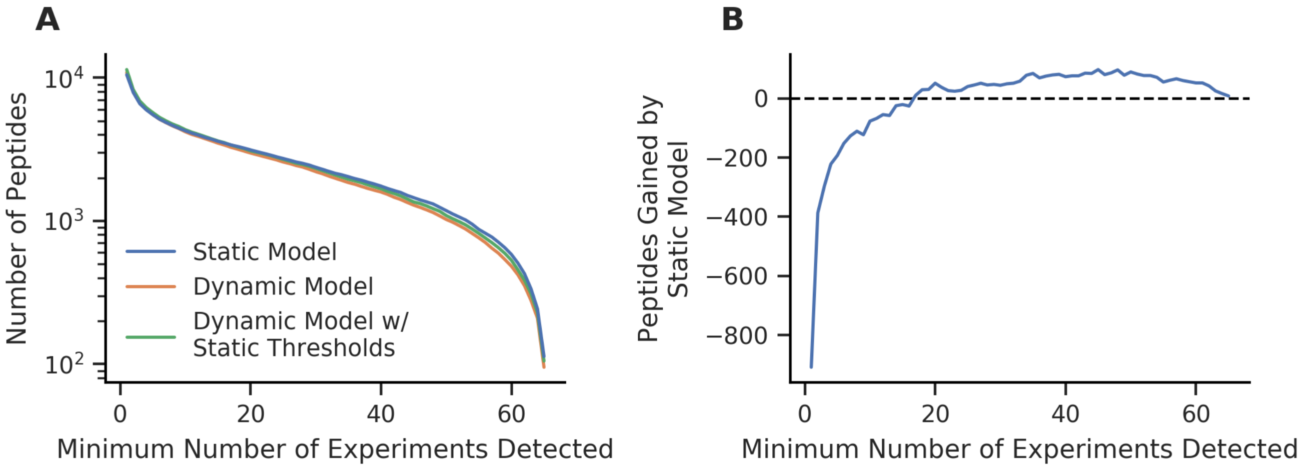


Supplementary Figure 2: The static Percolator model increases the number of peptides detected consistently across experiments. (A) The static model yielded fewer total unique peptides, but increased the number of peptides detected across many experiments in comparison to the Percolator's dynamically trained models. (B) When the dynamic model results were allowed the same number of peptides as the static model, the static model still detects more peptides that are present across high numbers of experiments.
